# Fluorescent reporter plasmids for single-cell and bulk-level composition assays in *E. faecalis*

**DOI:** 10.1101/797936

**Authors:** Kelsey M. Hallinen, Keanu A. Guardiola-Flores, Kevin B. Wood

**Author notes:** Email: K. Wood,.

## Abstract

Fluorescent reporters are an important tool for monitoring dynamics of bacterial populations at the single cell and community level. While there are a large range of reporter constructs available–particularly for common model organisms like *E. coli*–fewer options exist for other species, including *E. faecalis*, a gram-positive opportunistic pathogen. To expand the potential toolkit available for *E. faecalis*, we modified a previously developed reporter plasmid (pBSU101) to express one of nine different fluorescent reporters and confirmed that all constructs exhibited detectable fluorescence in single *E. faecalis* cells and mixed biofilm communities. To identify promising constructs for bulk-level experiments, we then measured the fluorescence spectra from *E. faecalis* populations in microwell plate (liquid) cultures during different growth phases. Cultures showed density- and reporter-specific variations in fluorescent signal, though spectral signatures of all reporters become clear in late-exponential and stationary-phase populations. Based on these results, we identified six pairs of reporters that can be combined with simple spectral unmixing to accurately estimate population composition in 2-strain mixtures at or near stationary phase. This approach offers a simple and scalable method for selection and competition experiments in simple two-species populations. Finally, we modified the construct to express codon-optimized variants of blue (BFP) and red (RFP) reporters and show that they lead to increased fluorescence in exponentially growing cells. As a whole, the results inform the scope of application of different reporters and identify both single reporters and reporter pairs that are promising for fluorescence-based assays at bulk and single-cell levels in *E. faecalis*.

## INTRODUCTION

*Enterococcus faecalis* is an opportunistic bacterial pathogen that is commonly found in the mammalian gut microbiome (1) and has been implicated in a range of (often drug-resistant) infections, including bacteremia, endocarditis, and wound infections (2, 3, 4, 5, 6, 7, 8). *E. faecalis* communities are characterized by a confluence of intercellular interactions, including communication via two-component signaling (9, 10, 11), pheromone-induced quorum sensing (12, 13), chromosomal transfer (14), extracellular electronic transfer (15), fratricide (16, 17), and phage-mediated gene transfer (18), leading to potentially complex population dynamics. Furthermore, the presence of antibiotics may induce rich ecological and evolutionary dynamics in *E. faecalis* (19, 20), making them an important model species for understanding the emergence and spread of drug resistance (21).

Fluorescent and bioluminescent reporters offer a straightforward method for monitoring growth dynamics of bacterial populations. For *E. faecalis* specifically, a number of different reporter constructs have used, offering several options for tracking dynamics–including growth, gene expression, and horizontal gene transfer–in single species populations (22, 23, 24, 25, 26, 27, 28, 29, 30). Unfortunately, simultaneous tracking of multiple populations remains challenging because of the limited spectral diversity of *E. faecalis* constructs and the relatively poor performance of fluorescent reporters–often optimized for common model species like *E. coli*– in low-GC gram-positive species (31, 32). In addition, *E. faecalis* cultures may exhibit decreased oxygen levels and low pH (33), potentially limiting the utility of some reporters. More general limitations to fluorescence include protein oligomerization and photobleaching (34, 35), obstacles that can often be addressed by optimizing proteins to a specific cellular context (36).

In this work, we set out to expand the fluorescent reporter toolkit available for the study of *E. faecalis* communities. To do so, we modified a reporter plasmid (pBSU101)– originally developed for fluorescent labelling of gram-positive bacteria for *in vivo* applications (23)–to constitutively express one of nine different fluorescent reporters, including variants of blue, red, green, and yellow fluorescent proteins. Because it was not clear, a priori, which reporters would be most effective in standard *E. faecalis* culture conditions, we chose a wide range of proteins from the commercially available ProteinPaintbox^®^ as well mTagBFP2 (BFP1) (37) and the native EGFP (GFP) contained in the original plasmid (23) (see Table 1). All constructs exhibited detectable fluorescence in single *E. faecalis* cells and mixed biofilm communities. For *E. faecalis* populations grown in microwell plate (liquid) cultures, we find that fluorescence signal in low density populations (OD600<0.05) cannot typically be distinguished from background signal of color-free controls, but spectral signatures of the different reporters become increasingly clear as density increases, particularly for late-exponential and stationary-phase populations. Based on these results, we identified six pairs of reporters that can be combined with simple spectral unmixing to accurately estimate population composition in 2-strain mixtures at or near stationary phase. Finally, we modified the construct to express codon-optimized variants of blue (BFP) and red (RFP) reporters and show that they lead to increased fluorescence in exponentially growing cells. As a whole, these results clarify the potential scope of application of different reporter constructs for bulk and microscopy-based experiments with *E. faecalis*.

**TABLE 1.** Fluorescent reporters used in this study.

| Nickname | Color | Origin |
| --- | --- | --- |
| GFP | EGFP | pBSU101 |
| BFP1 | mTagBFP2 | Addgene (RRID: Addgene_34632) |
| BFP2 | Optimized mTagBFP2 | GeneArt® Thermo Fisher Scientific® |
| PP-CFP | Cindy Lou CFP® | ATUM ProteinPaintbox® |
| PP-YFP1 | Yeti YFP® | ATUM ProteinPaintbox® |
| PP-YFP2 | Cratchit YFP® | ATUM ProteinPaintbox® |
| PP-GFP1 | Comet GFP® | ATUM ProteinPaintbox® |
| PP-GFP2 | Dasher GFP® | ATUM ProteinPaintbox® |
| PP-RFP1 | Rudolph RFP® | ATUM ProteinPaintbox® |
| PP-RFP2 | Optimized Rudolph RFP® | ATUM ProteinPaintbox® |
| PP-RFP3 | Fresno RFP® | ATUM ProteinPaintbox® |

## RESULTS

### Nine-reporter fluorescent library based on modified pBSU101 plasmid

To investigate the potential of different fluorescent proteins for monitoring *E. faecalis* populations, we modified plasmid pBSU101, which was originally derived from the shuttle vector pAT28 (39) and contains pUC and pAm*β* 1 (native to *E. faecalis* (38)) origins of replication for *E.coli* and a wide range of gram-positive organisms, respectively (23). We exchanged the native GFP from the pBSU101 plasmid with one of eight different fluorescent protein sequences (Figure 1, Table 1; see Methods). We also recircularized a color-free plasmid backbone, which serves as a fluorescence-free control. All modifications were confirmed via Sanger Sequencing, and final constructs were transformed into the *E. faecalis* strain OG1RF (40) (see Methods) to create nine different reporter strains.

**FIG 1.**
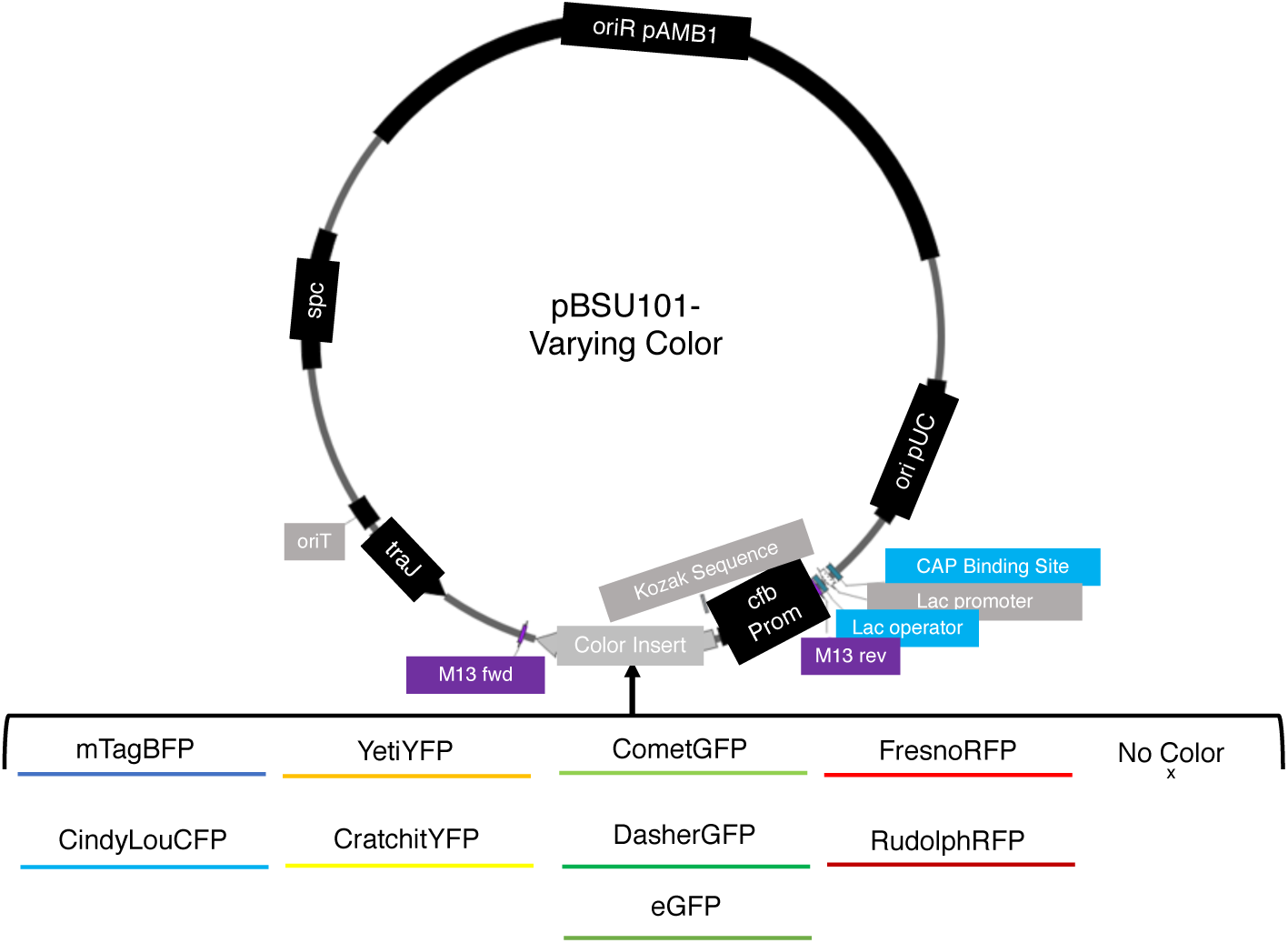
Schematic of reporter library based on pBSU101 plasmid. The reporter library was created with the vector backbone of pBSU101 (23), which contains a Spectinomycin resistance gene (*spc*) and constitutively expresses a fluorescent color behind a CAMP-factor gene (*cfb*) promoter from *S. agalactiae*. The vector also contains pUC and pAm*β* 1 (native to *E. faecalis* (38)) origins of replication for *E.coli* and a wide range of gram-positive organisms, respectively (23). The original *egfp* sequence was replaced with that for one of eight fluorescent reporters (See Table 1) as well as a recircularized color-free control.

### Imaging of single cells confirms visible fluorescence for each construct

To evaluate the functional utility of the different constructs, we grew cells overnight from single colonies, plated a diluted sample on a glass coverslip following several hours of growth, and imaged single cells using confocal microscopy (Figure 2). At this stage, our goal is merely to verify expression of the different reporters via fluorescence in the expected spectral range. The level of fluorescence varies across constructs, an effect that is at least partially attributable to the differing excitation protocols associated with each construct (see Methods). Nevertheless, we were able to confirm fluorescence in the appropriate spectral range for each reporter construct; by contrast, color-free control plasmids did not exhibit detectable fluorescence above background levels.

**FIG 2.**
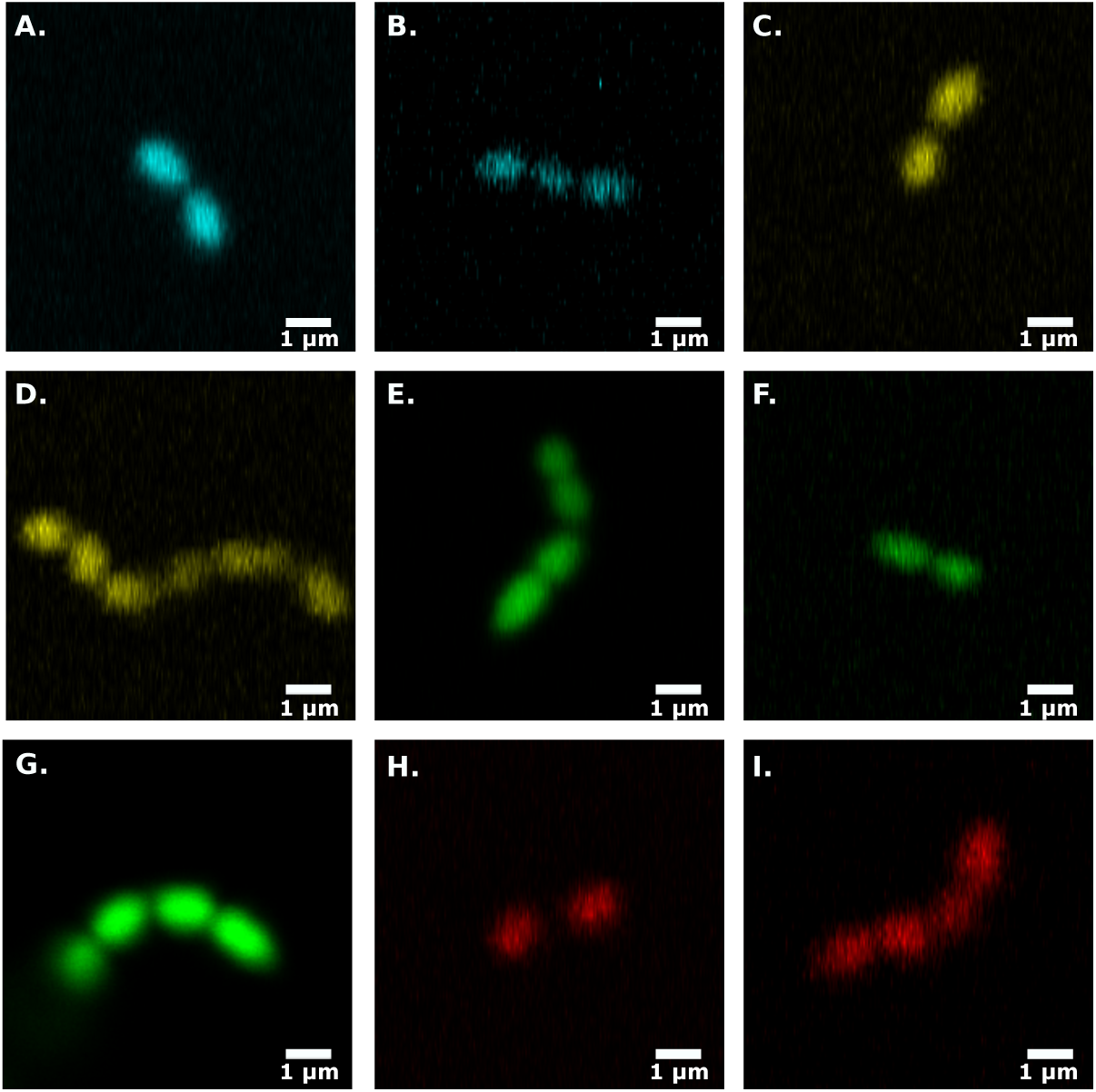
Single cell images of cells labeled with each different fluorescent plasmid. Confocal images of *E. faecalis* cells (see Methods) transformed with modified pBSU101 plasmid expressing BFP1 **(A)**, PP-CFP **(B)**, PP-YFP2 **(C)**, PP-YFP1 **(D)**, GFP **(E)**, PP-GFP1 **(F)**, PP-GFP2 **(G)**, PP-RFP1 **(H)**, or PP-RFP3 **(I)**.

Next, we wanted to evaluate the utility of the different reporter constructs for multi-color measurements in *E. faecalis* biofilms. Many fluorescent proteins, including those from the ProteinPaintbox^®^, require well-oxygenated environments for optimal maturation of their chromophore (23). However, biofilms are heterogeneous structures with regions of reduced O_2_ concentrations (41, 42). It is therefore not clear which–if any–of the fluorescent constructs will be visible under the non-optimal conditions represented by a biofilm. To answer this question, we grew 24-hour biofilms on glass coverslips starting from inocula comprised of two or three differentially labeled (but otherwise identical) *E. faecalis* populations (see Methods). We are able to clearly distinguish multiple subpopulations in 2D slices (Figure 3), indicating that these constructs may be useful for biofilm imaging of multicolor communities.

**FIG 3.**
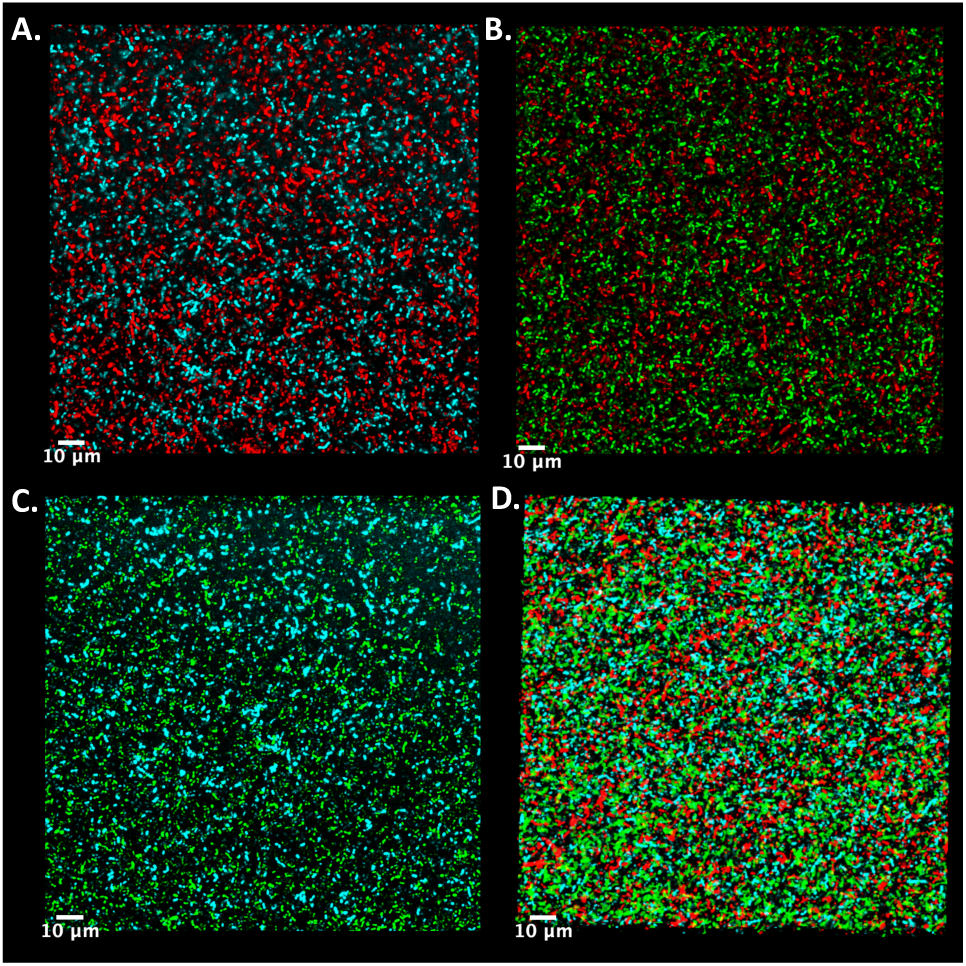
Representative 2D confocal images of two-color biofilms. Biofilms containing differentially labeled subpopulations were grown on coverslips overnight (24 hours) in a 6-well microplate and imaged using a Zeiss LSM700 confocal microscope (Methods). Biofilms contained mixtures of cells expressing BFP1 or PP-RFP1 **(A)**, PP-GFP2 or PP-RFP1 **(B)**, BFP1 or PP-GFP2 **(C)**, and BFP1 or PP-GFP2 or PP-RFP1 **(D)**.

### Distinct spectral features of different reporters are measurable in liquid cultures grown in microwell plates

Single cell images indicate that the reporter constructs lead to measurable levels of fluorescence that vary significantly across constructs. To quantitatively assess these differences, and to evaluate the feasibility of using these constructs for bulk-level experiments in liquid cultures, we grew *E. faecalis* populations–each transformed with a specific reporter construct–in commonly used *E. faecalis* media (Brain Heart Infusion, BHI) supplemented with Spectinomycin (120 *µ*g/mL) in 96-well microplates and measured time series of both optical density and the emission spectra of each population over a range of wavelengths surrounding the expected emission peaks for each reporter (Methods). To assess fluorescence signal, we define relative intensity (*I*_*r*_) as the fractional increase in fluorescence relative to that of a control population harboring a color-free variant of pBSU101 at the same density (*I*_*r*_ ≡ (*I* − *I*_0_)/*I*_0_, where *I* is the fluorescence intensity of the measured population and *I*_0_ that of the color free control, which we refer to as background).

We found spectral peaks in the expected wavelength range for each of the nine constructs, though the relative intensity of the peaks varied substantially (Figure 4; see Methods). In general, intensity increased as populations approached late exponential and stationary phase. For some constructs (e.g. PP-RFP3 and PP-GFP1), fluorescence intensity exceeded background (color-free control) by only 20-30 percent, even at the highest densities, while low density populations exhibited spectra indistinguishable from background. In other cases–for example, PP-YFP2 and especially PP-GFP2–spectral signatures were evident even in mid-exponential phase, while fluorescence in highdensity populations exceeded background by 200 percent or more.

**FIG 4.**
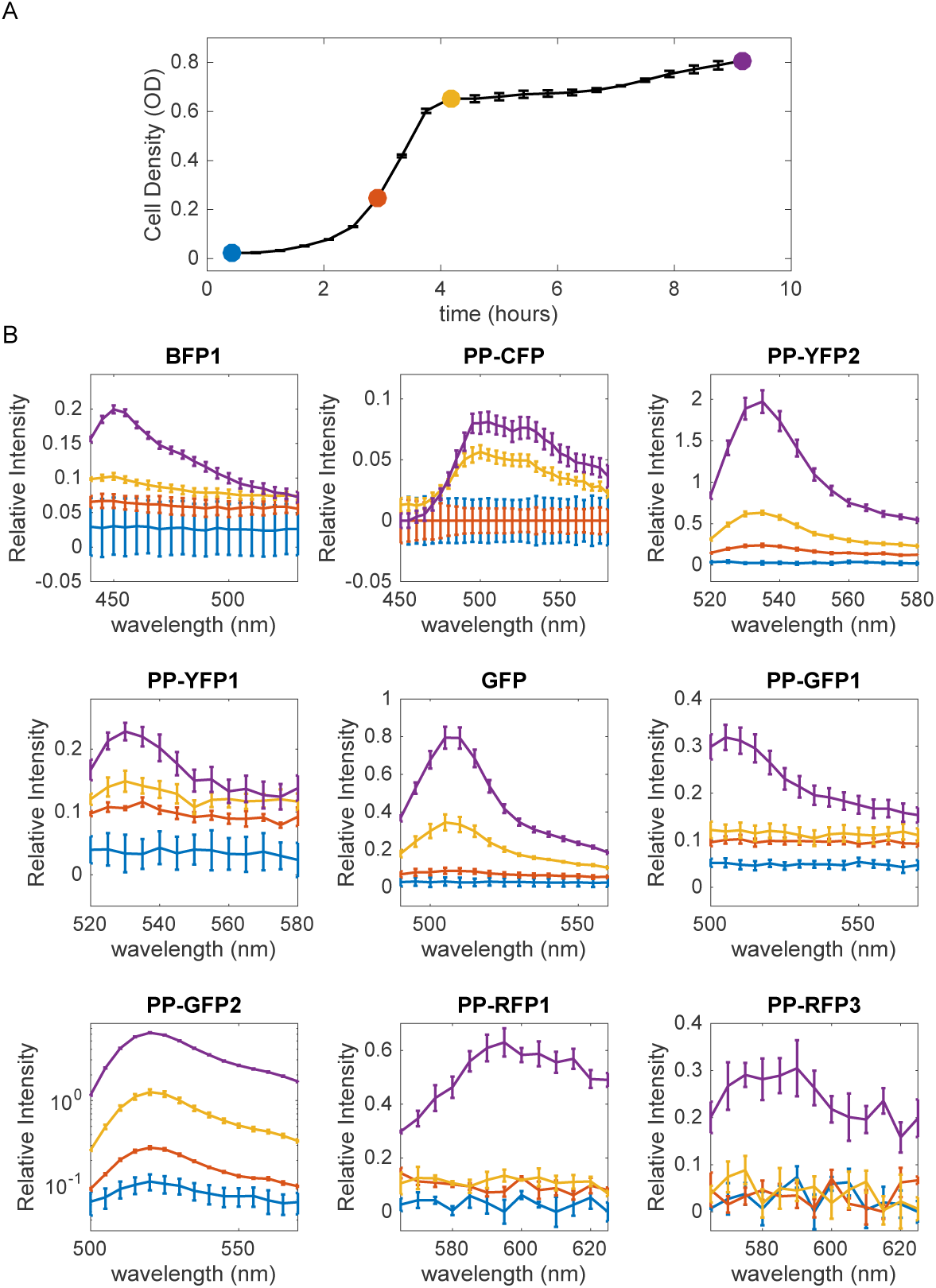
Fluorescence of bulk cultures varies significantly across growth phases and fluorescent reporters. **A.** Growth curve (OD600 time series) for *E. faecalis*. Circles represent densities at which color spectra were taken (part B). **B.** Emission spectra for populations transformed with one of nine reporter plasmids, with fluorescent reporter indicated in title (see Table I; see also Methods for excitation and emission settings). The relative intensity (*I*_*r*_) is defined as the fractional increase in fluorescence relative to that of a control population harboring a color-free variant of pBSU101 at the same density (*I*_*r*_ ≡ (*I* − *I*_0_)/*I*_0_, where *I* is the fluorescence intensity of the measured population and *I*_0_ that of the color free control). Note that vertical axes for the different panels have different limits. In particular, PP-GFP2 is shown on a log-scale to highlight a broader range of signal.

### Spectral unmixing to estimate population composition in simple mixtures

Constructs from the reporter library exhibit a range of different–and at times, non-overlapping–emission spectra with distinct spectral features. We therefore asked whether it was possible to quantitatively estimate the composition of simple populations containing two subpopulations, each with a different reporter plasmid. The emission spectra (Figure 4) suggest that some reporters–for example, pairs of YFP and GFP variants–would be difficult to unmix because of spectral overlap. On the other hand, other pairs are characterized by widely separated peaks and may therefore be good candidates for unmixing. While composition of fluorescent populations is often estimated using more sophisticated approaches–such as flow cytometry (FACS) (43, 44) or microscopy (45)–an assay based on a standard microplate reader would be convenient and could be scaled to achieve relatively high-throughput measurements.

As proof of principle, we created mixed communities of two populations–each harboring a different reporter construct– at different ratios. We chose six pairs of reporters (BFP1:PP-GFP2, BFP1:PP-YFP2, PP-CFP:PP-GFP2, PP-CFP: PP-RFP1,PP-GFP2:PP-RFP1, and PP-RFP1:PP-YFP2) where emission peaks were relatively well-separated. The mixtures were made from growing populations in spectinomycin-supplemented BHI in late exponential phase / early stationary phase. We then measured emission spectra for a limited span of wavelengths around the expected emission peaks for each of the reporters in the mixture (Methods). Note that for each pair of reporters, this protocol involves two different emission scans: one following excitation appropriate for the first reporter and one following excitation for the second reporter. We also repeated the same procedure for homogeneous populations harboring each of the individual reporters.

Using these measurements–and assuming that spectral features from different sub-populations combine linearly to produce spectra of the resulting mixture–we estimated from the measured spectra the population composition for each of the population mixtures (Figure 5). For all six reporter pairs, the demixing procedure allowed us to accurately and reliably estimate population composition of all 3 mixtures (Figure 5). These six pairs offer a range of color options, suggesting this demixing procedure may be easily adapted to different experimental setups.

**FIG 5.**
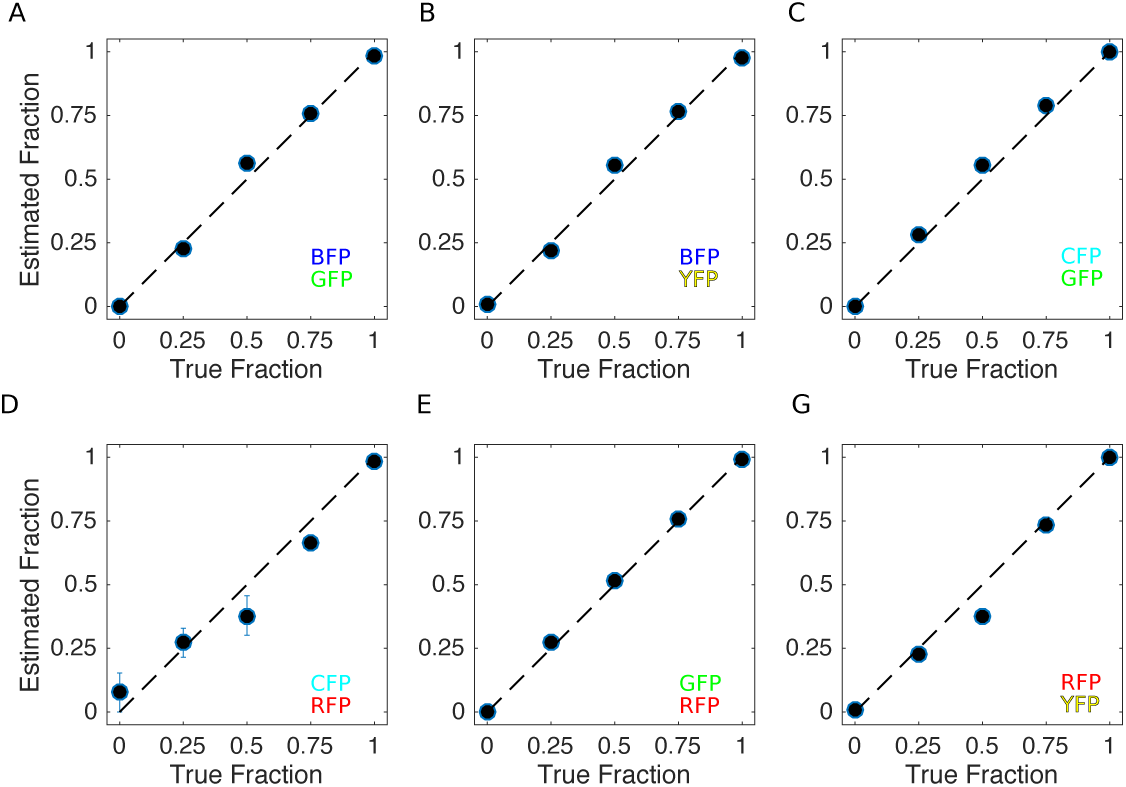
Composition of simple mixtures can be estimated using spectral unmixing in bulk cultures. Estimated population composition (vertical) vs true population composition (horizontal) for mixtures containing six pairs of reporter-labeled subpopulations: **A.** BFP1 and PP-GFP2, **B.** BFP1 and PP-YFP2, **C.** PP-CFP and PP-GFP2, **D.** PP-CFP and PP-RFP1, **E.** PP-GFP2 and PP-RFP1, **F.** PP-RFP1 and PP-YFP2. To create each mix, high density (near stationary phase) single-color populations were mixed in appropriate ratios and aliquoted in triplicate into individual wells of a 96-well microplate. For mixtures, circles represent the mean of three ummixing calculations, with error bars ± standard error of the mean. For single colors-0% or 100% one color, circles represent the mean of two unmixing calculations with error bars ± standard error of the mean.

### Codon optimization leads to slight increases in fluorescent intensity

Our re-sults identify several reporters in the green / yellow spectral range that are excellent candidates for single-cell and population level studies. On the other hand, BFP and RFP variants generally showed lower levels of relative fluorescence. In an effort to increase fluorescence signal from these constructs, we designed sequence variants of BFP1 and PP-RFP1 (BFP2 and PP-RFP2, respectively) that were codon optimized for expression in *E. faecalis* (Methods) and introduced those sequences into the pBSU101 construct. In bulk cultures, we found that populations transformed with BFP2 (optimized) and populations with BFP1 (non-optimized) exhibited similar levels of fluorescence in early exponential and late stationary phase. However, the populations with the codon-optimized construct showed higher fluorescence at both mid and late-exponential phases (Figure 6). By contrast, the PP-RFP2 (optimized) construct showed measurable increases in fluorescence only in late stages of growth, particularly in late stationary phase. We also found that single cells with BFP2 and PP-RFP2 taken from exponentially growing populations showed an increase in mean fluorescence relative to cells with the non-optimized counterparts (Figure 2).

**FIG 6.**
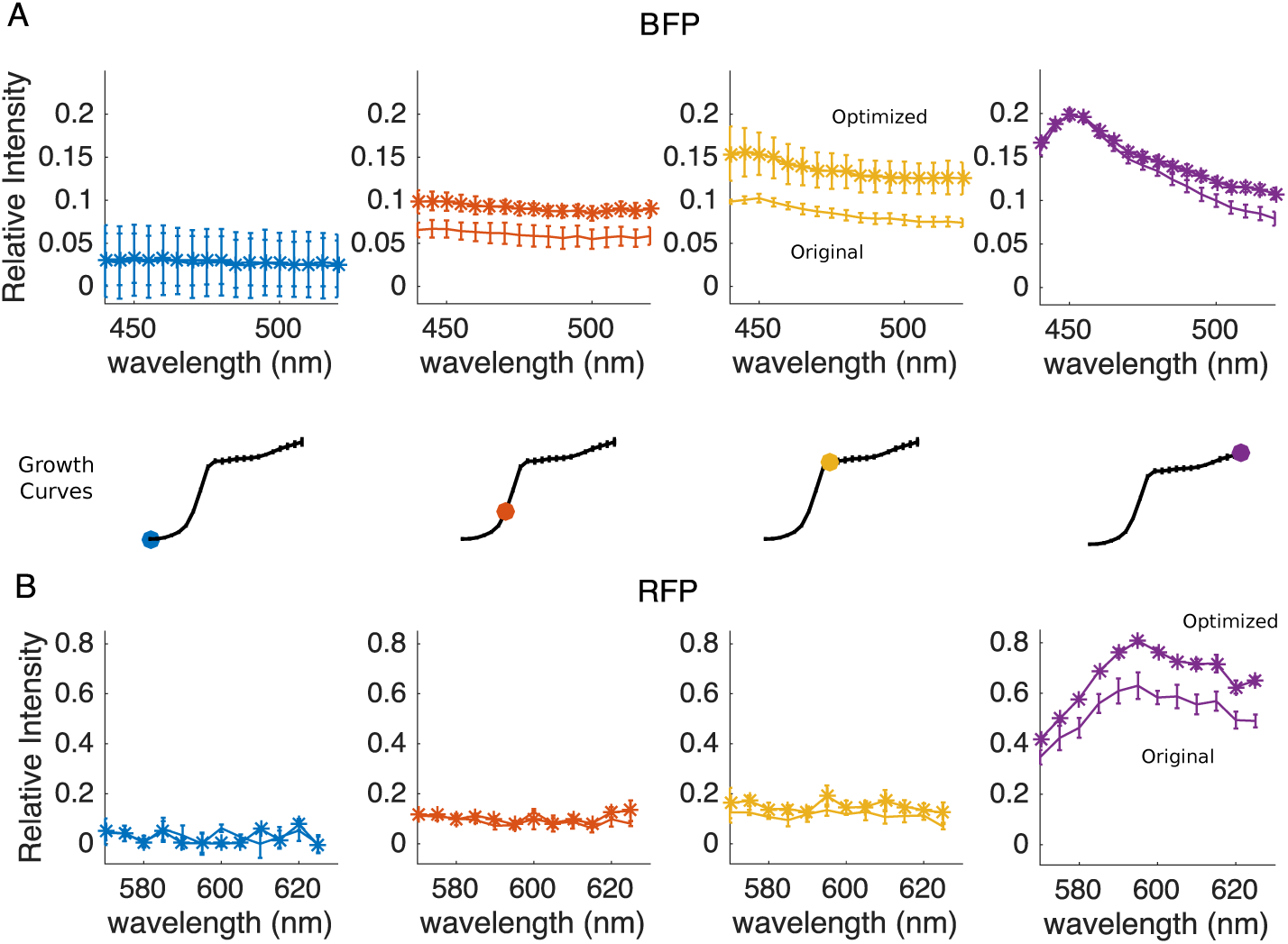
Codon optimized fluorescent reporters lead to population density-dependent increases in fluorescence intensity. The fluorescent spectra of BFP1 vs BFP2 **(A)** and PP-RFP1 vs PP-RFP2 **(B)** measured on a multimodal plate reader (Methods) at different stage of growth, ranging from early exponential phase (left, blue) to stationary phase (right, purple). Insets indicate specific regions of the growth curves where spectra were taken. Stars (non-stars) represent codon optimized (original) fluorescent reporters. Error bars are ± one standard error of the mean.

**FIG 7.**
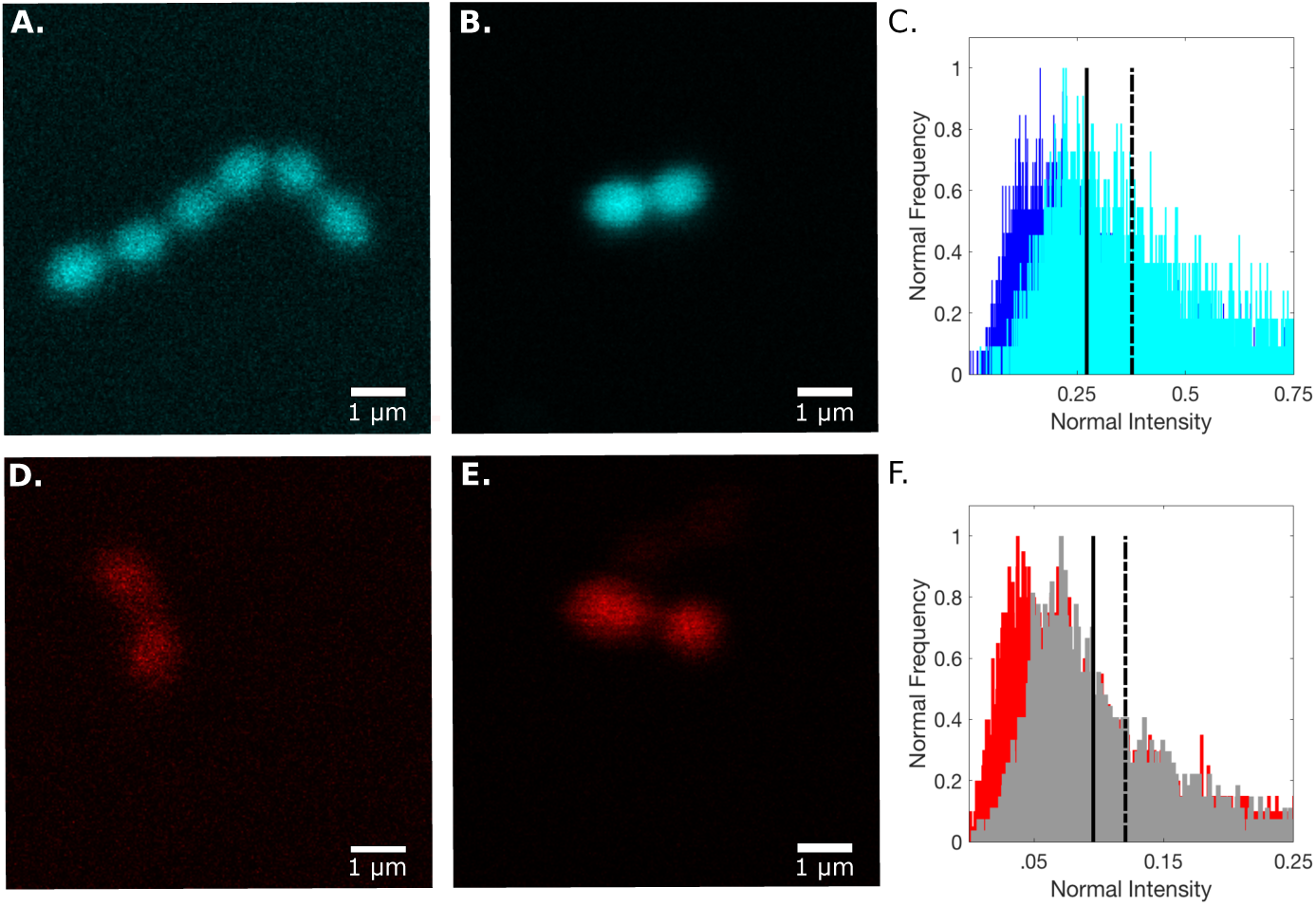
Codon optimized reporter constructs increase fluorescent signal in individual *E. faecalis* cells. Example confocal images of single cells expressing BFP1 **(A)**, BFP2 **(B)**, PP-RFP1 **(D)**, or PP-RFP2 **(E)** during exponential growth (Methods). Right panels: histogram of normalized intensity of populations expressing either BFP1 (dark blue) or BFP2 (cyan) **(C)** or either PP-RFP1 (red) or PP-RFP2 (gray) **(F)**. Solid (dashed) lines represent the mean intensity averaged over all cells for the non-optimized (optimized) reporter. Error bars are ± standard error of the mean across three replicates.

## DISCUSSION

In this work, we modified a previously developed reporter plasmid (pBSU101) to constitutively express one of nine fluorescent reporters with different spectral properties and evaluated the potential utility of these constructs for tracking *E. faecalis* populations. We observed detectable fluorescence in single *E. faecalis* cells and mixed biofilm communities, while populations grown in bulk cultures showed density- and reporter-specific variations in fluorescent signal. Some reporters–for example, reporter PP-GFP2–showed readily detectable spectral signatures, even in low density populations, while others (PP-RFP3) could only be reliably measured in high-density stationary phase populations. Based on these results, we identified six pairs of reporters that can be combined with simple spectral unmixing to accurately estimate population composition in 2-strain mixtures at or near stationary phase. Finally, we created constructs with codon-optimized variants of blue (BFP) and red (RFP) reporters and show that they lead to increased fluorescence in exponentially growing cells.

It is important to keep in mind several limitation of this work. As noted previously, fluorescence signal under both *in vitro* and *in vivo* conditions may depend on a number of factors, such as oxygen concentration and pH, which are not well controlled in our experiments. Our goal was to evaluate performance of these constructs under standard and convenient (*in vitro*) conditions, but different environments–particularly those *in vivo*–may lead to different results. At the same time, further optimization of *in vitro* conditions–for example, the use of fluorescent-friendly media with low background–will likely improve signal for at least some of the reporters considered here.

We have limited our estimates of population composition to cultures at high densities (nearing stationary phase), where fluorescent signal was particularly bright, and to simple pairwise mixtures. While it may be possible to achieve similar levels of accuracy with mixtures containing the brightest reporters (e.g. PP-GFP2) at lower densities or in larger mixtures, other reporter combinations are sure to be limited to the simplest scenarios. It is also not clear whether this approach can be used for dynamic measurements in growing populations, where density is changing over time. Optimizing media–or alternatively, resupending cells in PBS or similar media–may improve composition estimates, making the method more widely applicable. But even in the absence of these extensions, the method provides an easily scalable option for competition assays in high-density populations, such as those involving daily “endpoint” measures of composition (43, 44, 46).

In summary, we have constructed a plasmid-based fluorescence reporter library for *E. faecalis* and quantified the performance of different constructs in single-cell and bulk measurmenets. Our results may be useful for researchers looking to select reporter proteins for different applications, as we identify both single reporters (e.g. PP-GFP2) and reporter pairs that are promising for fluorescence-based assays and also develop codon-optimized variants of BFP and RFP with improved fluorescent signal under some conditions. Finally, the simple spectral unmixing method presented here provides a convenient assay for estimating population composition in standard microwell plates, without the need for sophisticated approaches based on flow cytometry or single-cell microscopy.

## METHODS

### Bacterial Strains, Media, and Growth Conditions

All experiments were performed in strain OG1RF, a fully sequenced *E. faecalis* oral isolate (40). Overnight cultures were grown at 37°C in sterile REMEL™ BHI Broth–Brain Heart Infusion (37g in 1L demineralized water) following isolation of a single colony from BHI Agar plates–Fisher BioReagents™ Agar Powder/Flakes (15g/L). Media and plates were supplemented with Spectinomycin (120 µg/ml) (Spec120) (Spectinomycin sulfate; MP Biomedicals).

### Amplification of sequences for fluorescent reporter proteins

Sequences for seven fluorescent proteins (PP-CFP, PP-GFP1, PP-GFP2,PP-RFP1, and PP-RFP3) (see Table 1) were PCR-amplified from IP-Free© Fluorescent ProteinPaintbox™-*E. coli* (47, 48)). The sequence for BFP1 was similarly amplified from plasmid pBAD-mTagBFP2 (RRID:Addgene_34632, (37)). In both cases, PCR amplification of the relevant regions was performed using Thermoscientific™ Phire™ Hot Start II DNA Polymerase on plasmids isolated from stock cultures with the QIAprep^®^ Spin Miniprep Kit followed by linearization with appropriate restriction enzymes (NEB).

### Codon optimization of PP-RFP1 and BFP1

The sequences PP-RFP1 and BFP1 were codon optimized for expression in *E. faecalis*. Synthesis of the optimized BFP1 sequence (BFP2) was performed by GeneArt^®^ (Thermo Fisher Scientific^®^) and optimized PP-RFP1 (PP-RFP2) was done by ATUM. Sequences were codon-optimized based on codon usage tables available for *E. faecalis* strain V583.

### Construction of fluorescence reporter library based on pBSU101 plasmid

The fluorescent library was constructed by swapping the native *egfp* sequence in pBSU101– whose expression is driven by the constitutive promoter of *cfb* (23)–with the sequence for one of the eight different fluorescent proteins described above (see Table 1). The cloning was performed using standard Gibson Assembly (GA) methods (49). Primers used for PCR and Gibson Assembly for the 8 fluorescent proteins as well as those for the pBSU101 backbone are shown Tables 2 and 3. Table 4 shows the primers used for PCR and GA of the optimized color sequences. Table 4 also has the primers for the recircularized, no color plasmid.

**TABLE 2.**
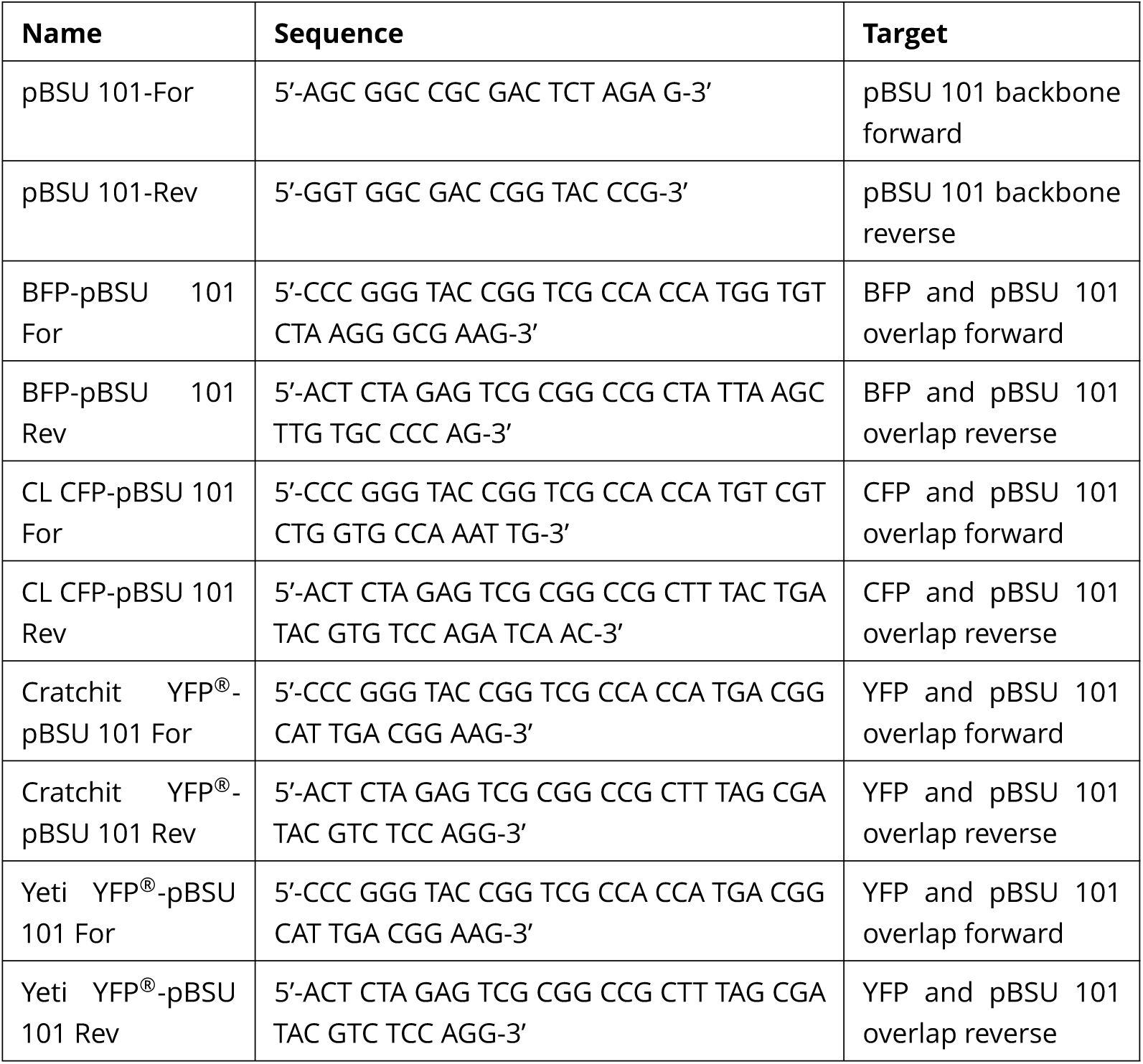
Primers used for PCR and Gibson Assembly of the 8 different fluorescent library plasmids. Cindy Lou CF^®^: CL CFP. Backbone primers for used for all the colors (optimized and non-optimized) are also listed here.

**TABLE 3.**
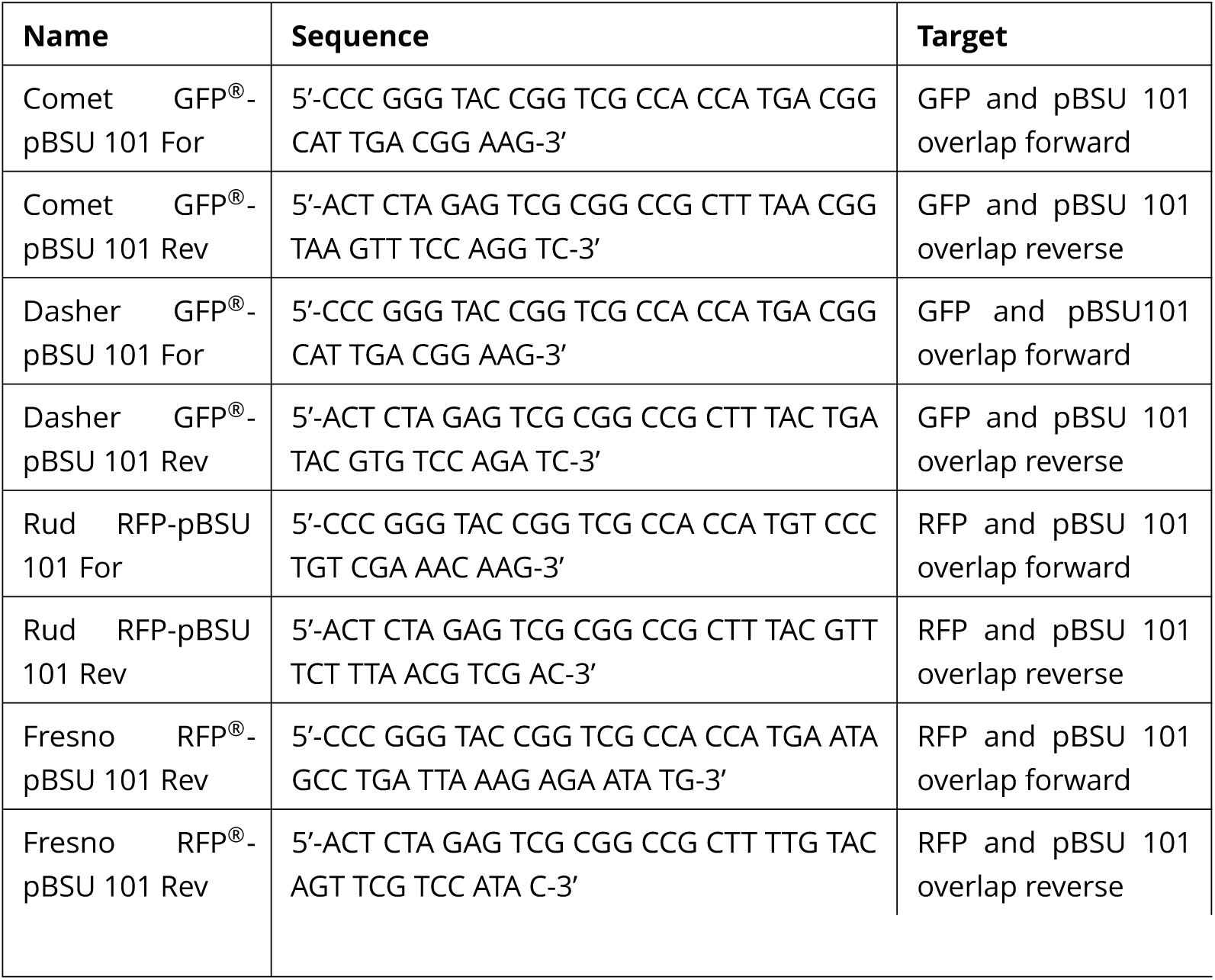
Primers used for PCR and Gibson Assembly of the 8 different fluorescent library plasmids, continued. Rudolph RFP^®^: Rud RFP.

**TABLE 4.**
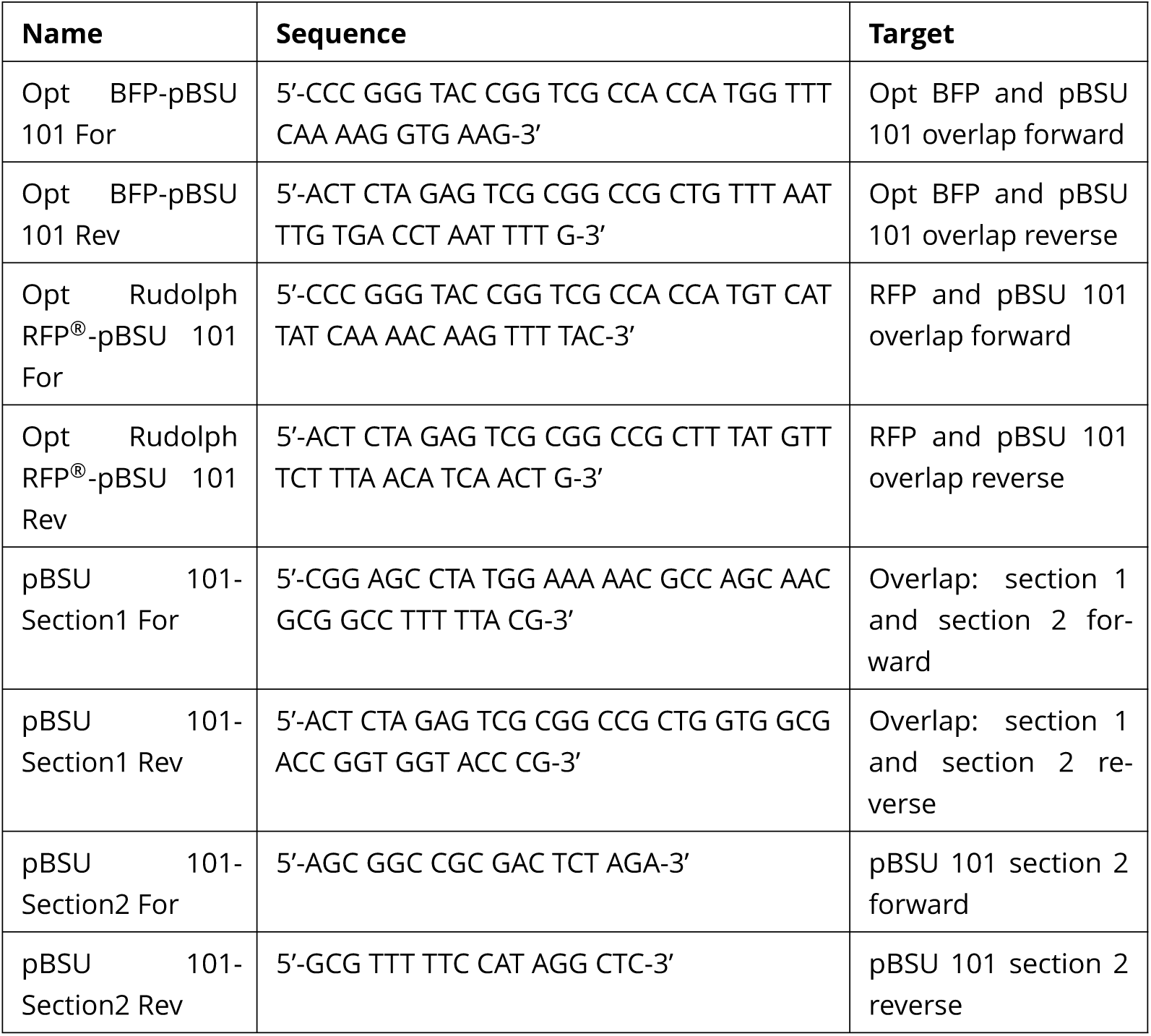
Primers used for PCR and Gibson Assembly of the optimized and no color plasmids. For the no color plasmid, PCR was used to create the backbone in two separate pieces and Gibson Assembled back together, without the color sequence.

Following construction of the different reporter constructs, the plasmids were originally transformed into the high efficiency cloning strain NEB^®^ 5-*α* Competent *E. coli* via heat shock transformations (50). Following successful transformations, plasmids were miniprepped from the NEB^®^ 5-*α* Competent *E. coli*, sequence verified, and transformed into the *E. faecalis* strain OG1RF, with electroporation (51).

### Plate reader experiments on bulk populations

Color spectra were measured in 96-well microplates using an Enspire multimodal plate reader. Overnight cultures of cell populations harboring each of the reporter constructs, including the no color control plasmid, were diluted to an optical density (OD600) of 0.01 at 600 nm. Fluorescence and OD600 were measured every 45 minutes from each well until the cells reached stationary state. The excitation and emission used for each color are shown below in Table 4.

### Estimating population composition with spectral unmixing

Mixed populations comprised of two sub-populations (each with a different reporter construct) were created by diluting cells in late exponential / early stationary state to a common OD600 and then mixing at designated ratios. Experiments were run immediately after mixing to avoid changes in composition due to growth. Emission spectra for the mixed populations were then measured using parameters in Tabel 6. For each 2-color mixed population, two scans were taken, one for each reporter construct.

To estimate population composition, we created single-color intensity vectors *s*_*i*_ for each homogeneous (single color) population *i*. Each vector had 12 components; the first six correspond to the (abbreviated) emission spectrum using parameters associated with the the first color, the second six to the (abbreviated) emission spectrum using parameters associated with the second color. The abbreviated emission spectrum is defined as the emission at a pre-determined six-point region surrounding the expected peak for each color (see Table 6). In an ideal experiment with no spectral overlap between colors, the six components of *s*_*i*_ corresponding to the reporter present in the population would be nonzero, while all other components would approach zero.

**TABLE 5.** Excitation and Emissions used for the plate reader full spectra. Emission spectra were taken in the range listed with a step size of 5 nm.

| Color | Excitation (nm) | Emission (nm) |
| --- | --- | --- |
| mTagBFP2 | 401 | 440-530 |
| CindyLou CFP® | 400 | 450-580 |
| Yeti YFP® | 500 | 520-580 |
| Cratchit YFP® | 500 | 520-580 |
| Comet GFP® | 480 | 500-570 |
| Dasher GFP® | 480 | 500-570 |
| EGFP | 470 | 490-560 |
| Rudolph RFP® | 545 | 565-625 |
| Fresno RFP® | 545 | 565-625 |

**TABLE 6.** Excitation and Emission wavelengths used for spectral unmixing experiments. Emissions spectra were taken at a total of 6 equally spaced wavelengths spanning the range above, which corresponds approximately to the peak for each reporter.

| Color | Excitation (nm) | Emission (nm) |
| --- | --- | --- |
| mTagBFP2 | 401 | 440-465 |
| CindyLou CFP® | 400 | 495-520 |
| Cratchit YFP® | 500 | 525-550 |
| Dasher GFP® | 480 | 510-535 |
| Rudolph RFP® | 545 | 585-610 |

We then measured the color-intensity vector, *m*, for the mixed population whose composition we wished to estimate. If fluorescent intensity were proportional to the size of each sub-population, the intensity vector *m* could be approximated as a linear combination of the vectors *s*_*i*_ corresponding to each individual color, with the *N* weighting coefficients *c*_*i*_ corresponding to the relative size of each of the *N* sub-populations.

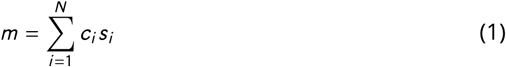

In our experiments, *N* = 2 and normalization (*c*_1_ + *c*_2_ = 1) reduces the number of free parameters to one. To estimate this parameter, we use simple linear regression (52).

### Confocal Microscopy of single cells and biofilms

Overnight cultures were diluted 1:5 in fresh BHI media supplemented with Spectinomycin (120 *µ*g/mL) and grown for 30 minutes before imaging with a LSM700 confocal microscope (Zeiss), 40X (1.4 N.A) oil objective and a pinhole of 1.2 microns. Table 7 shows the different excitation and emission protocols used.

**TABLE 7.** Excitation laser and emission spectra used for the confocal microscopy. These values are based on pre-loaded protocols in the Zeiss LSM700 software that correspond to fluorescent proteins with similar spectral properties.

| Color | Excitation Laser | Laser Power (mW) | Laser Attenuation (%) | Protocol |
| --- | --- | --- | --- | --- |
| mTagBFP2 | 405 nm | 5 | 5.0 | mBFP |
| CindyLou CFP® | 405 nm | 5 | 5.0 | mBFP |
| Yeti YFP® | 488 nm | 10 | 1.0 | Lucifer Yellow |
| Cratchit YFP® | 488nm | 10 | 1.0 | Lucifer Yellow |
| Comet GFP® | 405 nm | 5 | 5.0 | EGFP |
| Dasher GFP® | 488 nm | 10 | 0.5 | EGFP |
| EGFP | 488 nm | 10 | 0.5 | EGFP |
| Rudolph RFP® | 555 nm | 10 | 5.0 | mStrawberry |
| Fresno RFP® | 555 nm | 10 | 5.0 | mStrawberry |

To compare fluorescent signal in single cells with optimized vs non-optimized reporters, we imaged single cells at different time points in the growth process. Overnight cultures were diluted 1:100 in fresh BHI with Spectinomycin (120 *µ*g/mL), grown for 1.5 hours, and then imaged every hour (by plating a small volume onto a coverslip) for 15 hours. Boundaries of each cell were selected from each image using ImageJ, and the pixel intensity within each cell boundary was measured in MATLAB^®^ to determine intensity per cell at the different growth stages. Pixel intensities are represented as 16-bit integers. We normalize their intensities to a maximum value of 50,000. Frequencies are normalized to the highest frequency of pixel intensity values within the range of 0 to 50,000. Normalization of the frequencies was done for optimized and non-optimized distributions independently. In addition to generating histograms for the range of intensities, we also averaged each set of pixel intensities to obtain the mean intensity for each optimized and non-optimized distributions.

Mixed biofilms comprised of 2 or 3 differentially labeled sub-populations were grown from overnight (single color) cultures that were diluted 1:100 and then mixed in equal ratios to create two or three-color mixed populations. The mixtures were then grown on glass coverslips in 6-well plates overnight, which were removed after 24 hours and directly imaged.

